# Prediction of range expansion and estimation of dispersal routes of water deer (*Hydropotes inermis*) in the transboundary region between China, the Russian Far East and the Korean Peninsula

**DOI:** 10.1101/2022.02.16.480706

**Authors:** Ying Li, Yuxi Peng, Hailong Li, Weihong Zhu, Yury Darman, Dong Kun Lee, Tianming Wang, Gleb Sedash, Puneet Pandey, Amaël Borzée, Hang Lee, Yongwon Mo

**Affiliations:** Tiger and Leopard Conservation Fund in Korea (KTLCF), and Research Institute for Veterinary Science, College of Veterinary Medicine, Seoul National University, Seoul 151-742, 08826, Republic of Korea; College of Geography and Ocean Science, Yanbian University, Hunchun, Jilin 133000, China; National Forestry and Grassland Administration Key Laboratory for Conservation Ecology in the Northeast Tiger and Leopard National Park, Beijing 100875, China; Amur branch of World-Wide Fund for Nature, Verkheportovaya St., 18A, Vladivostok, 690003, Russia; Department of Landscape Architecture and Rural System Engineering, Seoul National University, Seoul 151-742, 08826, Republic of Korea; College of Life Sciences, Beijing Normal University, Beijing 100875, China; Northeast Tiger and Leopard Biodiversity National Observation and Research Station, Beijing 100875, China; Land of the Leopard National Park, 100-letiya Vladivostoka 127, Vladivostok, 690068, Russia; Laboratory of Animal Behaviour and Conservation, College of Biology and the Environment, Nanjing Forestry University, Nanjing, People’s Republic of China; Department of Forest Resources and Landscape Architecture, College of Life and Applied Sciences, Yeungnam University, Gyeongsan, Republic of Korea

**Author notes:** ***Corresponding Author:*** Hang Lee, Tiger and Leopard Conservation Fund in Korea (KTLCF), and Research Institute for Veterinary Science, College of Veterinary Medicine, Seoul National University, Seoul 151-742, South Korea, Email address Yongwon Mo, Department of Forest Resources and Landscape Architecture, College of Life and Applied Sciences, Yeungnam University.

**Keywords:** Water deer, dispersal corridor, species distribution model, conservation planning, circuit theory

## Abstract

Global changes may direct species expansion away from their current range. When such an expansion occurs, and a species colonizes a new region, it is important to monitor the habitat used by the species and use the information for updated management strategies. Water deer is listed as Vulnerable species in IUCN Red List and restricted to east central China and the Korean Peninsula. Since 2017 water deer has expanded its range towards northeast China and the Russian Far East. Our research focuses on data collected in northeast China and the Russian Far East during 2017-2021, with the purpose of providing support for a better understanding of habitat use and provide conservation suggestions. We used MaxEnt to model species niche and distribution and predict habitat suitability for water deer and applied the circuitscape to determine possible dispersal routes for the species. There is good quality habitat for water deer in the boundary area of the Yalu and Tumen River estuaries between China, North Korea, and the Russian Far East, as well as the east and west regions of the Korean Peninsula. Elevation, distance to cropland and water sources, and presence of wetlands were the variables that positively contributed to modelling the suitable habitats. Two possible dispersal routes were determined using the circuit theory, one was across the area from North Korea to the downstream Tumen transboundary region, and the other was across North Korea to the boundary region in China and along the tiger national park in northern China. A series of protected areas in North Korea, China, and Russia may support the dispersal of water deer. The establishment of a Northeast Asia landscape conservation network would help establish monitoring and conservation planning at a broad scale, and this study provides an example for the need for such a network

## 1. Introduction

Herbivore animals play an important role in the ecosystem in terms of direct ecological interaction with animal predator and vegetation used for food. Herbivores also have indirect interactions with other species, such as insect biodiversity [1, 2]. Monitoring the habitat used by herbivores is as important part of the work needed for conserving the whole ecosystem. Human activity, climate change and population growth of the focal species may cause species to shift their original distribution. An important point for monitoring is that when a species first occupies a new region, it will show clearer habitat preferences compared to areas where it has long been established.

Water deer (*Hydropotes inermis*) is a species endemic to East China (Chinese water deer, *H. i. inermis*) and the Korean Peninsula (Korean water deer; *H. i. argyropus*). The Water deer is listed as ‘Vulnerable’ at a global level by the IUCN Red List [3]. The population of Chinese water deer is decreasing and occurring in a fragmented habitat. The main distribution area of the Korean water deer is the Korean Peninsula. In South Korea, water deer are described as a widespread species, as they occupy most of the possible habitats, including wetlands, grasslands, and forests [4]. In North Korea, water deer are listed as wildlife with economic value and have been provided government support for protection in the wild. Water deer populations were historically distributed in western North Korea, and this species was relocated three times to the east of North Korea in 1968 [5]. In 2005, the presence of water deer was reported from protected areas located in eastern North Korea, including Cheonbul Mountain Animal Reserve (천불산동물보호구) in South Hamgyeong Province and Daegak Mountain Animal Reserve (대각산동불보호구) in North Hwanghae Province [6]. Although there are no specific occurrence data or population descriptions of water deer in North Korea, combining the available references, it can be assumed that water deer are distributed in suitable habitats in both western and eastern North Korea [3, 6]. However, since 2019, water deer have been frequently reported in the boundary area between China, Russia, and the Democratic People’s Republic of Korea, where there are no earlier formal records of this species [7]. In the newly occupied areas in the lower Tumen River basin in China and in the Russian Far East, there are still many wild areas with relatively low human impact. Therefore, new individuals dispersing into the region may choose their preferred habitat. As herbivores, water deer contribute to ecosystem functions, and they can be a potential prey for big cats such as tigers and leopards in the national parks in the boundary area between China and Russia. Habitat use can provide information on the potential expanding trend and dispersal routes. Knowing the habitat of this species is essential for management and conservation of local wildlife and ecosystems. Water deer prefer wetlands, swamps, low lands, and grasslands, and do not avoid farmlands [8]. Along the Yalu and Tumen rivers, new water deer records are cropping up, as there seem to be large areas of suitable habitat along the riverbanks. As this deer species can quickly increase in population size [9], it may become an important mid-sized mammal species in newly occupied territories, which may also affect river and wetland ecosystems, and may also impact the distribution of other wildlife.

It is urgent to understand the habitat used by new expansions of population, predict the distribution of the expansions and the possible dispersal route. Species distribution model and landscape analysis can help provide the knowledge by considering species and their environmental variables. Species distribution modeling (SDM), also called environmental, bioclimatic, species niche, or habitat suitability modeling [10], can be used to assess the habitat use of a species. The results can reveal the factors that may influence the species, and thus may provide important references for management of the species [11]. SDM can also be applied to predict species dispersal under climate change scenarios, as well as to model the results of changes in land use [12]. Climate change data and land use prediction data are often used separately, but numerous unexpected biases may be included because of limitations on the knowledge of species population size or land cover connectivity [13, 14]. Maximum entropy (Maxent) is a widely used machine learning method for SDMs, and it was specifically developed for presence-only data to overcome the problems of small undesigned samples [15].The core idea is to take full account of the known information when inferring the distribution of unknown probabilities and treating the unknown information indiscriminately [16]. Habitat suitability level can also be predicted, thus enabling the calculation of difference in habitat and resource \between protected and non-protected areas [17]. Circuit theory is a recently developed technique that can quantify movement across a landscape and can be applied in many fields, including landscape ecology, population genetics, movement, and fire behavior [18]. This method is based on random walk and graph theories [19]. A circuit is usually defined as a network of nodes connected by resistors and is used to analyze graphs.

Our research focuses on understanding and clarifying the current expansion of water deer, as dispersing individuals are entering new territories. As a potential invasive species, it is imperative to understand the species habitat preferences and predict the range expansion in order to guide management strategies for monitoring and research in the newly occupied areas. We used a Maxent model to predict water deer habitat characteristics, and the circuit method to analyze the conservation area network in the Tumen transboundary region. The results provide an overall knowledge of the potential habitat and possible dispersal routes of water deer, which can be used for further monitoring and conservation strategies for wildlife on a larger scale.

## 2. Material and methods

### 2.1. Research overview and data processing

Our research area focused on the new expansion area including the southern regions of Liaoning, Jilin, and Heilongjiang provinces in China, the Primorye region in Russia, the whole of North Korea, and the northern part of South Korea (34.70317°N–49.008799°N, 117.65389°E– 139.26630°E). Ninety-nine occurrence records of water deer from the new expansion region were used for our habitat modelling (Figure 3). The occurrence data came from various sources, including ecological monitoring, camera trapping records, roadkill and hunting records, as well as published literature records [20-22] and the public database Global Biodiversity Information Facility (GBIF) [23]. To avoid overfitting when conducting the habitat model, we deleted the occurrence points within in 500 meters of each other, resulting in 75 data points used for the analyses [24].

**Figure 1.**
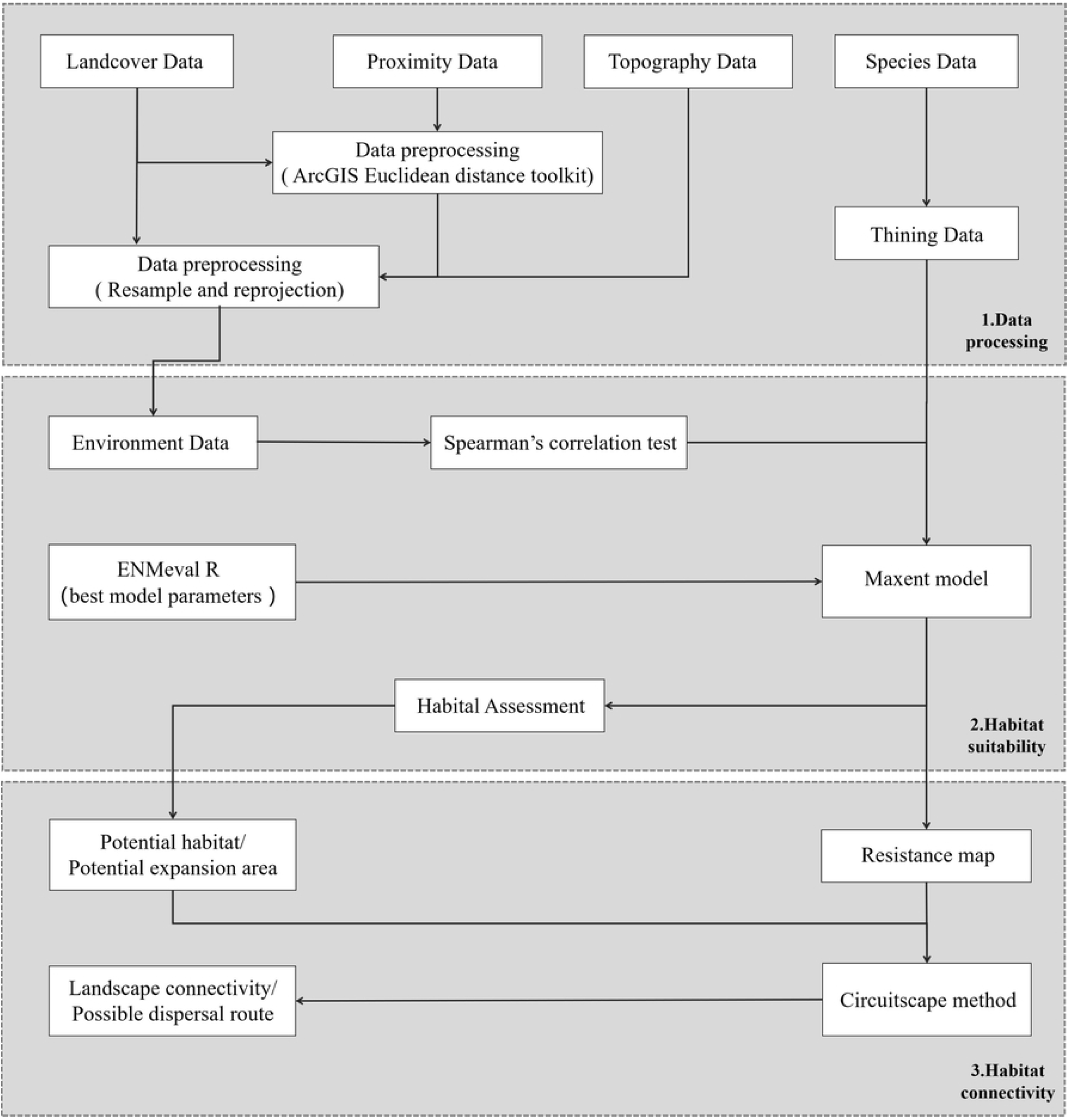
Flowchart of the framework used for water deer expansion modeling

**Figure 2.**
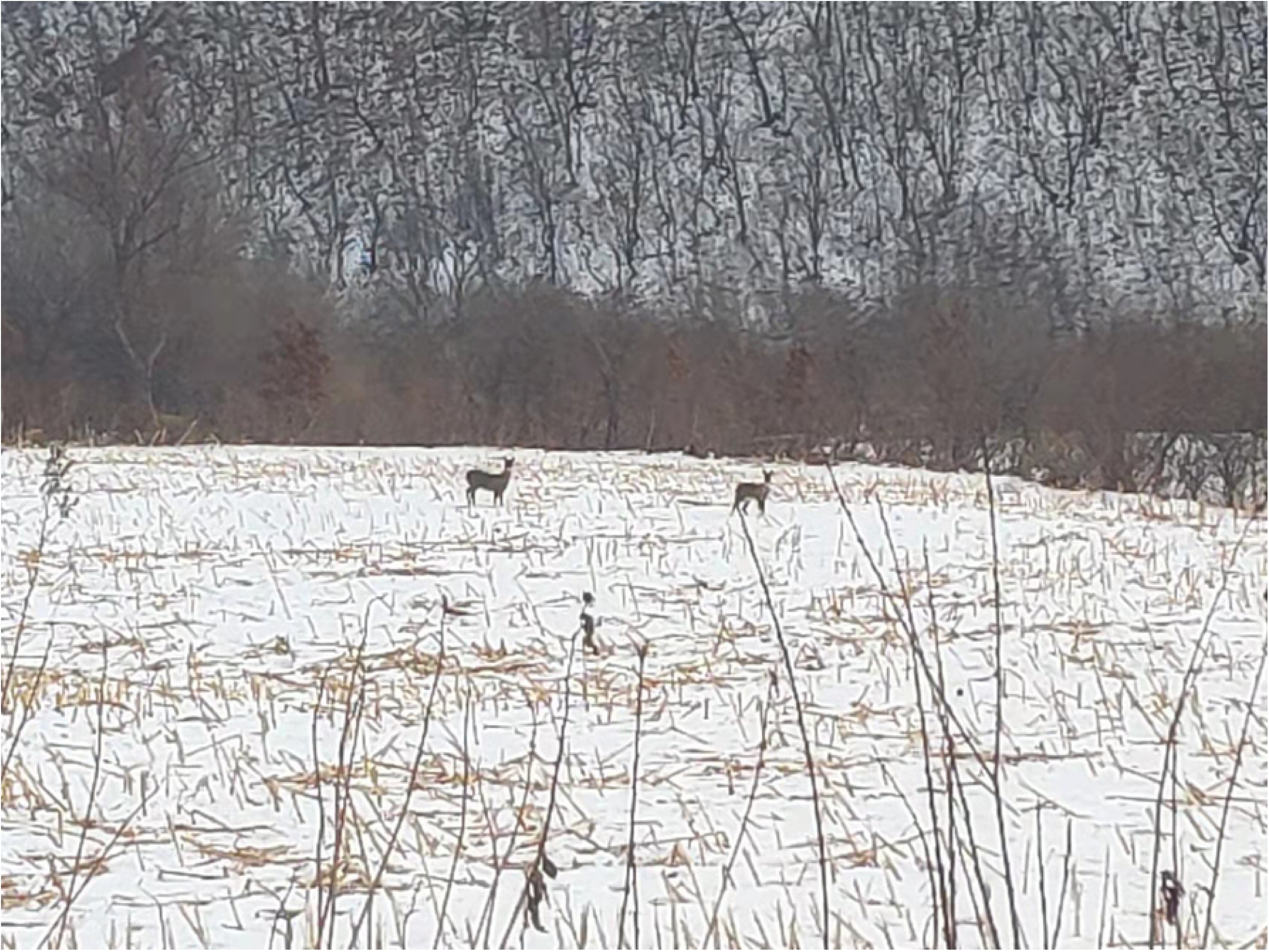
Forest edge with cropland and grassland habitat - the typical environment where water deer were spotted. Captured in the winter of 2019, Hunchun, northeast China.

**Figure 3.**
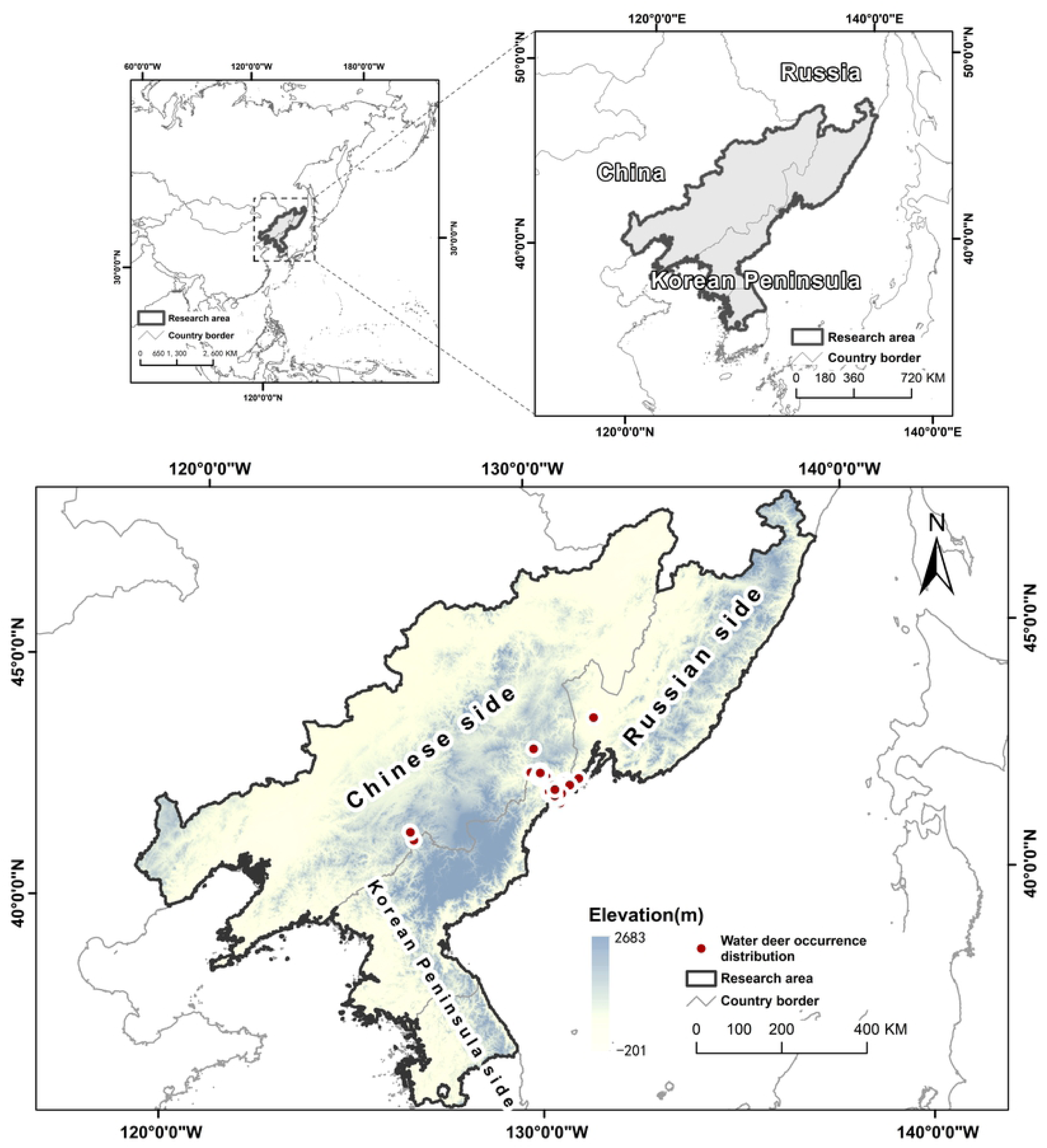
Locations of water deer occurrence used for species distribution modeling. We combined research data including survey data from camera trapping, regular monitoring data as well as data from public database and published references in the new expanded northeast China and the Russian Far East during 2019-2021 (n=102).

Considering the water deer ecology in southeast China and South Korea, seven main environmental variables (Table 1) were selected for habitat modelling including altitude, slope, aspect, distance to built-up areas, distance to water, distance to cropland and distance to roads [9, 25, 26]. The three topographic data utilized were altitude, slope, and aspect, which were processed from the SRTM 90-meter resolution World DEM database[27].

**Table 1.**
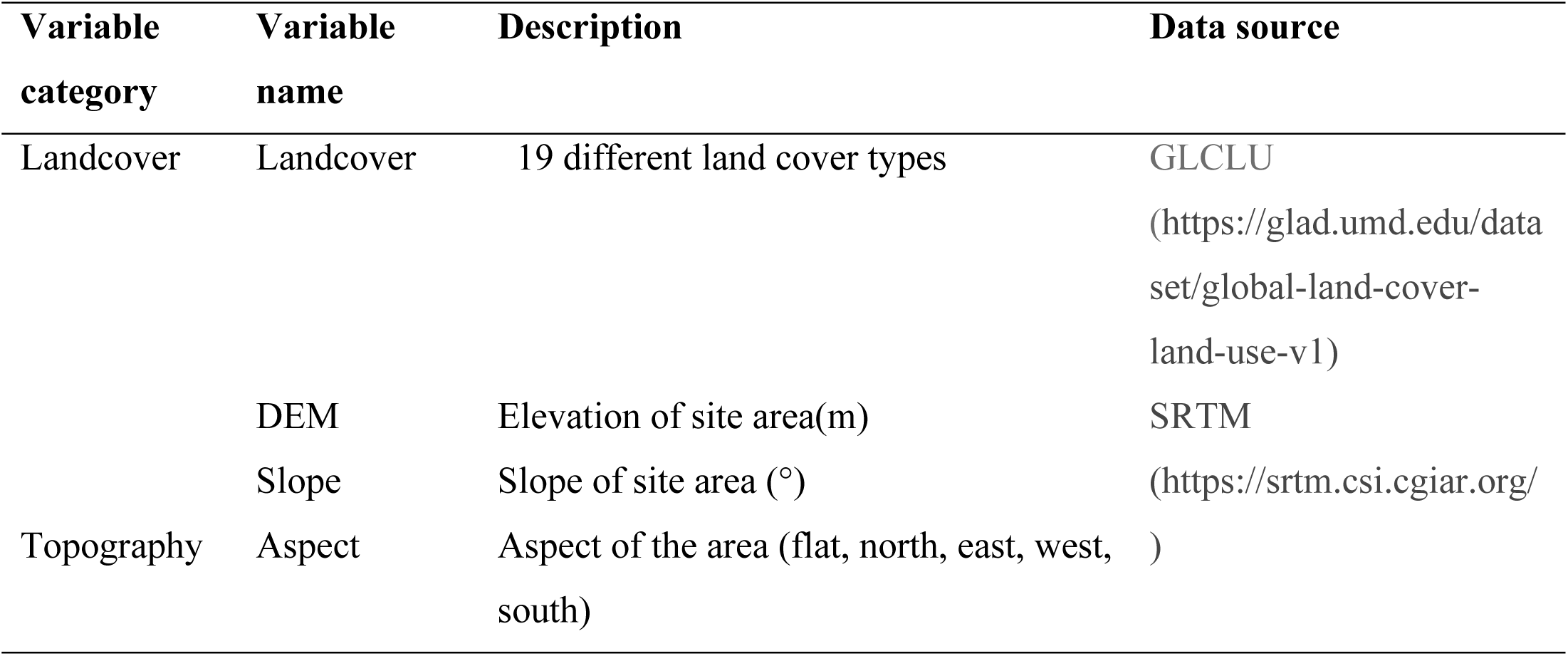
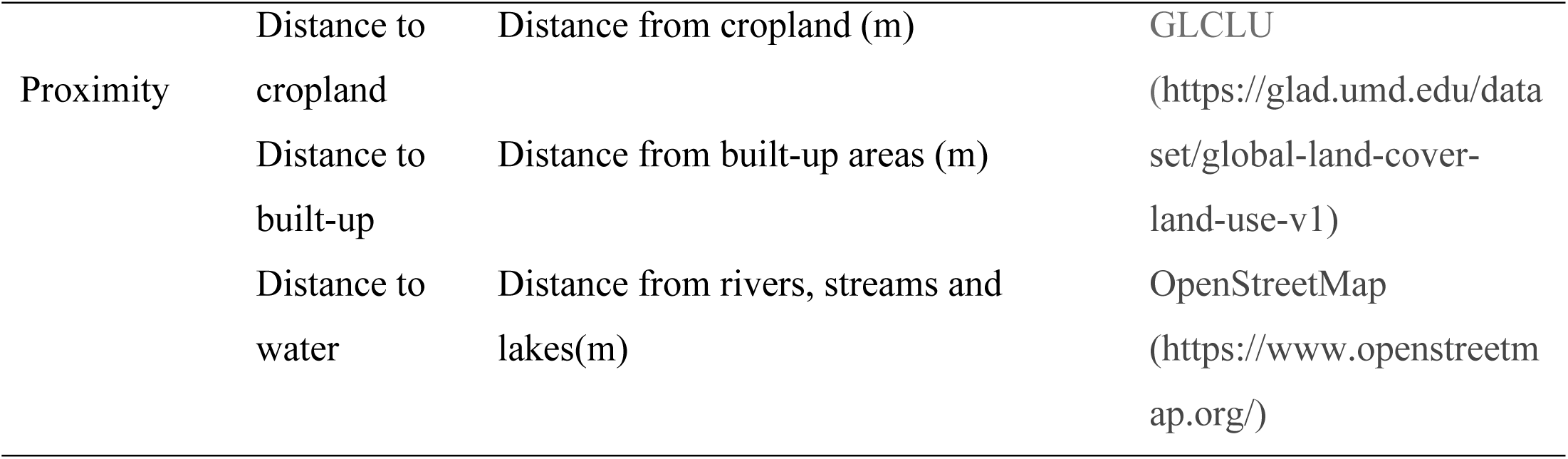
Water deer environmental variable

Altitude and slope variables were continuous data, and aspect was classified into five categories ranging from to 0–360 degrees: 1-flat (−1–0.0001 degrees), 2-north (315–360 degrees), 3-east (45–135 degrees), 4-south (135–225 degrees), and 5-west (225–315 degrees). Land cover data was processed from global land cover [28] and land use data from 2019 with 19 land cover types, namely desert, semi-arid, dense short vegetation, open tree cover, dense tree cover, tree cover gain, tree cover loss, salt pan, wetland sparse vegetation, wetland dense short vegetation, wetland open tree cover, wetland dense cover, wetland tree cover gain, wetland tree cover loss, ice, water, cropland, built-up area, and ocean [28]. Distances from built-up areas, cropland, and water were processed using the ArcGIS Euclidean distance toolkit, with data from land cover and the open street map database as sources [28, 29]. To obtain a high-quality predictions, and since strongly correlated environmental variables can lead to model overfitting, we conducted a Spearman’s correlation test using the IBM SPSS software version 26.0 [30] and selected seven uncorrelated environmental variables to estimate habitat suitability. The variables thus selected were aspect, slope, distance to water, distance to cropland, distance to built-up, elevation, and landcover. Considering the home range of water deer, we selected a 500 m resolution for the analyses [31], and we used the 2000 Korean Central Belt 2010 projection system.

### 2.2. Habitat suitability

In this study, we used Maxent 3.4.1 software [32, 33] to build the habitat model. We uploaded the seventy-five occurrence point data and the seven uncorrelated environmental variables to the software, and we set the output format as Logistic, using 25 percent species occurrence data as test data set. Maxent performance is closely related to both the regularization multiplier parameter and feature selection [34]. The ENMeval R package was applied to test the best model parameters by setting it to select the model with the most significant result, omission value less than 5%, and delta AICc value less than 2 [35]. The best model setting was selected using linear, quadratic, and threshold features together, with the regularization multiplier value set to 0.9. We built the model with 10,000 background points and 10 bootstrap replicates. A Jackknife analysis was used to assess the contribution of environmental variables. The model performance was evaluated using a threshold-independent area under the curve (AUC) of the receiver operating characteristic curve (ROC). The Jenks method was employed to determine habitat suitability level using ArcGIS 10.3 (ESRI, Redland, CA, USA).

### 2.3. Habitat connection

Based on Ohm’s law, voltage V is applied across a resistor, current I flows through it, and the current intensity depends on the voltage V and resistance R [36].In the application of circuit theory to landscape ecology, complex landscapes can be considered as the conductance, and the species randomly travelling to different directions can be considered as the random walker. Ecological features that are beneficial (positive) for the movement of the focal species, such as preferred land cover types, are given low resistance values, whereas negative features are given high resistance values.

Connectivity was analyzed for three areas (named as region 1, region 2 and region 3) with high-quality habitat, as defined using Maxent results. The newly occupied areas in the lower Tumen region were defined as target region TM1. The predicted high-value habitat in the west and east coasts of the Korean Peninsula, for which there are also records of water deer [3, 6], were set as regions NK2 and NK3 respectively [3, 6]. The three high quality habitat patches regions TM1, NK2 and NK3 were designated as nodes, and thus the target regions for the analyses on connectivity. The three patches were given specific values with value 1, 2, and 3. The background values were set as 0 and the layer was saved as ascii file.

For the background we had to define resistance surfaces to calculate the resistance between the nodes. The surface resistance can also be calculated from the Maxent modelling results [37], and we used the surface values derived from the Maxent results using the formula R = 101 − M, where R is the surface resistance and M is the Maxent value*100. Because Maxent values multiplied by 100 ranged from 0 to 100, the resistance surface values thus ranged from 1 to 101 [38]. The calculations were conducted in ArcGIS raster calculator. The resistance surface and node map were converted to the ascii data and uploaded to the Circuitscape ArcGIS toolkit to determine the connection map [39], using all settings as default.

## 3 Results

### 3.1. Habitat suitability

The model’s ROC curve had an AUC value of 0.935±0.014, indicating that our model could accurately simulate the relationship between the geographical distribution of water deer and the factors analyzed. The three most important variables selected for the model were elevation and distances to cropland and water body (Table 2). Elevation had the highest percentage of contribution to the model (59.6%), followed by distance to cropland (16.3%), land cover (10.8%), and distance to water sources (8.8%). Other variables contributed 4.4% overall to the model, which were slope (2.1%), followed by aspect (1.6%) and distance to built-up areas (0.7), indicating that these three variables were less important for the model compared to the other variables.

**Table 2.**
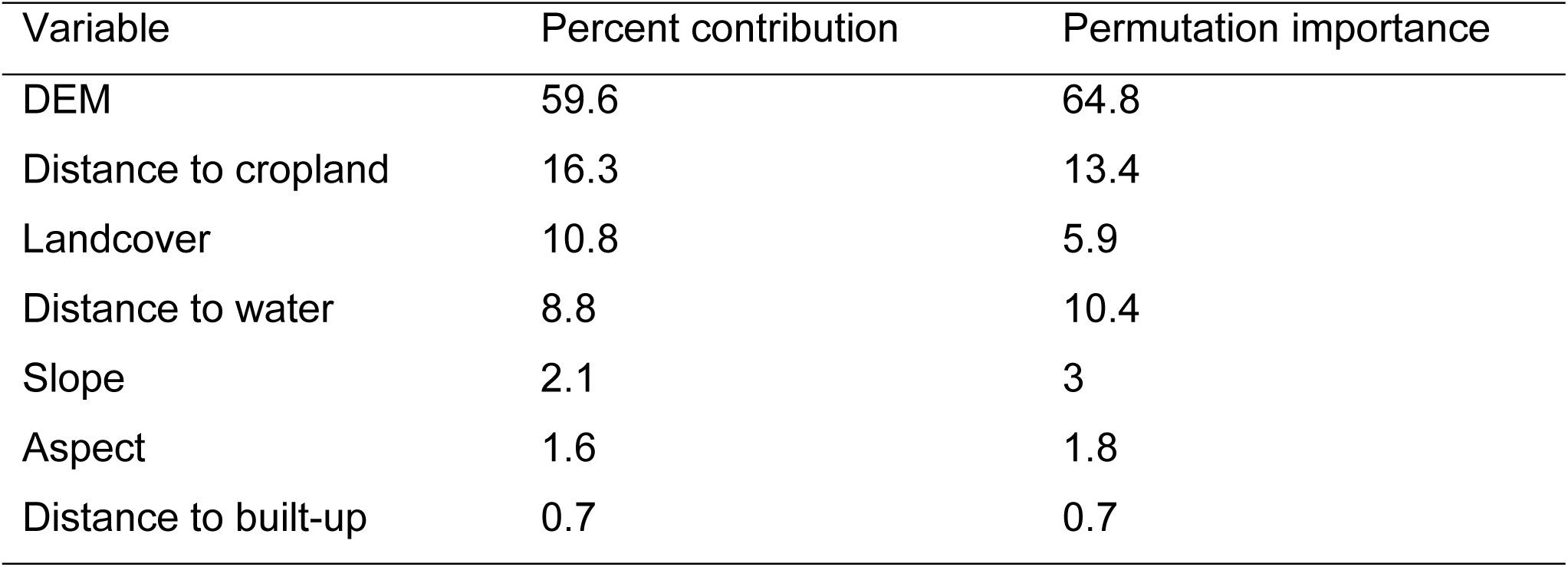
Environmental variable contribution percent and permutation importance used for water deer habitat suitability modelling. Based on 99 distribution locations in Northeast China and the Russian Far East

| Variable | Percent contribution | Permutation importance |
| --- | --- | --- |
| DEM | 59.6 | 64.8 |
| Distance to cropland | 16.3 | 13.4 |
| Landcover | 10.8 | 5.9 |
| Distance to water | 8.8 | 10.4 |
| Slope | 2.1 | 3 |
| Aspect | 1.6 | 1.8 |
| Distance to built-up | 0.7 | 0.7 |

Based on the response curves, from a logistic output value above 0.5, a higher probability of presence could be determined for each variable. Factors that contributed to the highest probability of water deer presence were elevation below 100 m, dense short vegetation, wetland with dense short vegetation and open tree cover, and distance below 1000 m from water sources and croplands.

The habitat with the highest suitability was mainly found in three areas. One comprised of the west coast of the Korean Peninsula, stretching into the southern Liaoning province in northeast China. The second was found along the east coast of the Korean Peninsula, ranging to the north of the Russian Far East coastal area. The third area comprised the area in which water deer have recently dispersed into the lower Tumen basin and stretched 740 km to the north along the Ussuri River.

### 3.2. Habitat connectivity

Region TM1 comprised the areas newly occupied by water deer, located in the lower Tumen River basin. Region NK2 was in the western part of North Korea, including North Pyongan province, the western part of South Pyongan province, the Pyongyang region, and the western parts of North and South Hwanghae provinces. Region NK3 included the southern part of South Hamgyong province and the northern area of Kangwon province in eastern North Korea. Our results show strong connectivity between regions NK2 and NK3, but only weak connectivity between these regions and region TM1. Two possible dispersion routes were estimated based on the landscape connectivity analysis. Route A connected the habitat between regions NK3 and TM1, whereas South Hamgyong Province, except the eastern area, connected to North Hamgyong province, through Ryanggang province and the border area between China and North Korea, and passes the northern area of North Hamgyong province to the north and connected to the region downstream of the Tumen River. Route B possibly passes north of South Pyongan province and South Hamgyong province to Chagang province and passes the border between China and North Korean south of the Baishan area and through Linjiang, Fusong, and south of Antu, Wangqing, and Hunchun before reaching region TM1 downstream of the Tumen River region and connecting region NK2 to region NK3 (Figure 6).

**Figure 4.**
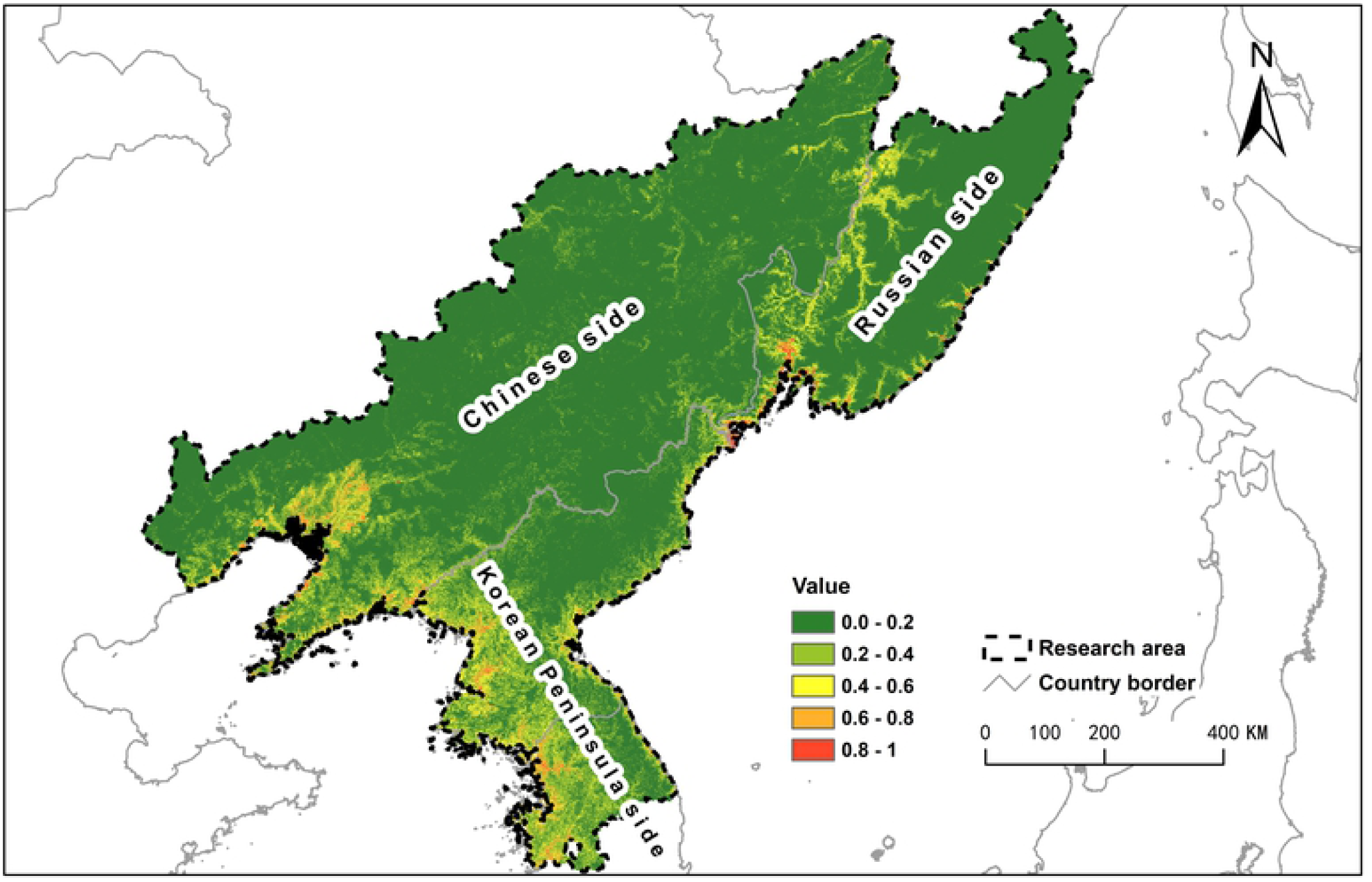
Maximum Entropy model of habitat suitability for water deer (AUC=0.939 ±0.011). The high suitability region (0.4-1) located along the Korean Peninsula’s east and west coasts towards northeast China and the Russian Far East.

**Figure 5.**
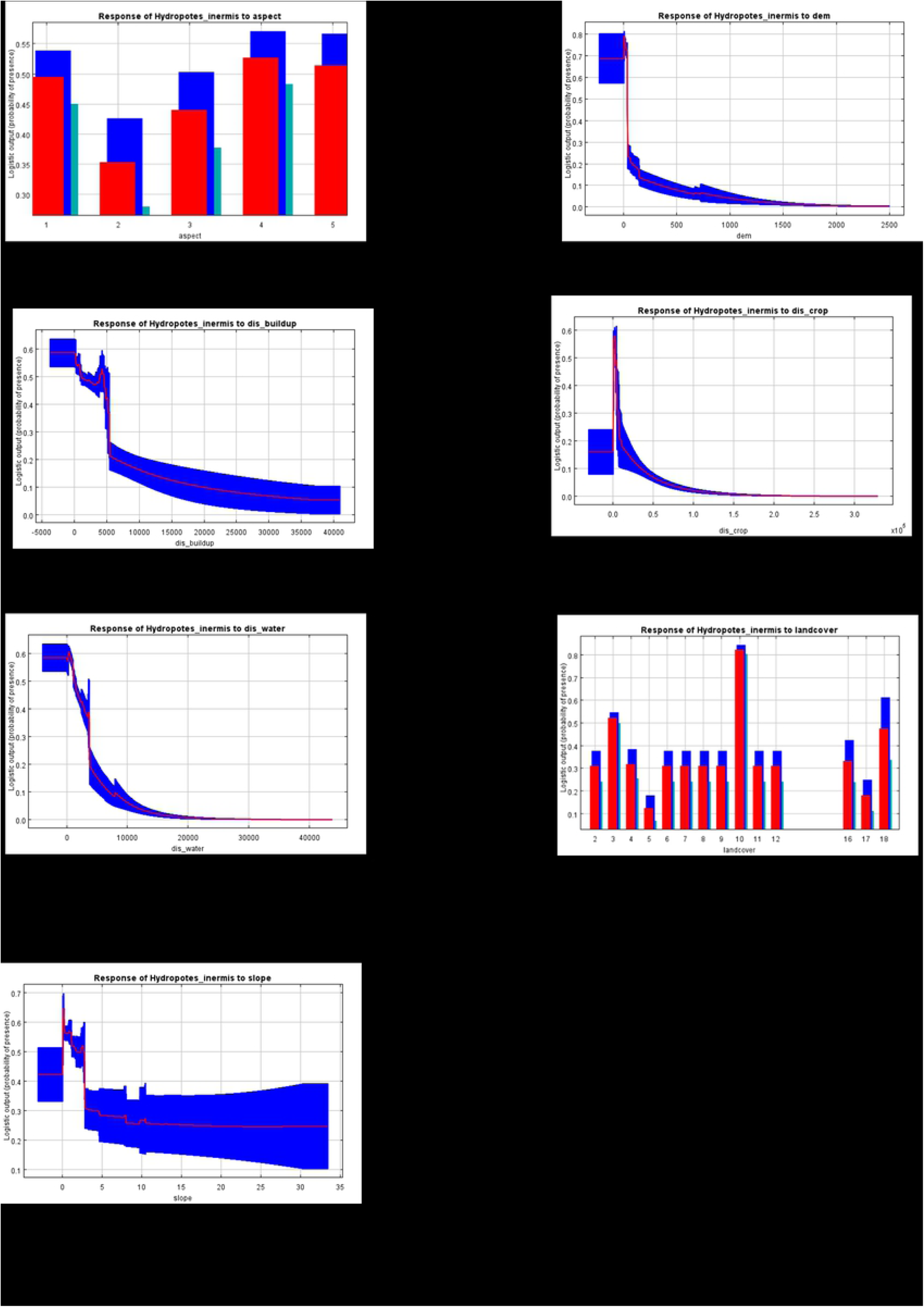
Response curves of water deer to environmental variables used for model prediction. (A:Aspect; B: DEM; C: Distance to built-up; D: Distance to cropland; E: Distance to water; F: Land cover type ; G:Slope)

**Figure 6.**
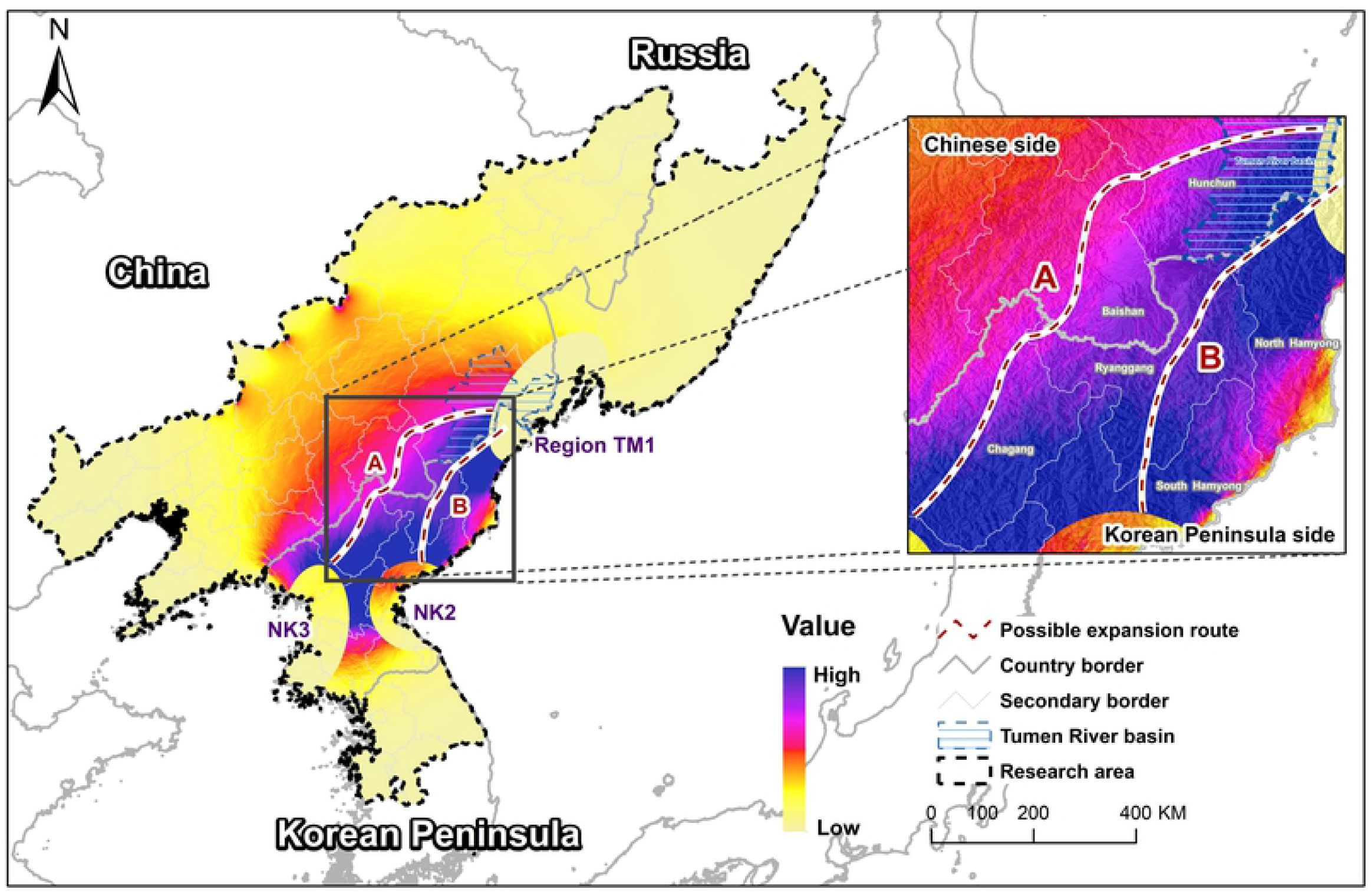
Possible dispersal routes of water deer. Suitable habitat patch region TM1, NK1 and NK2 were calculated the connections, two possible routes were estimated, route A cross North Korea into China boundary area reach the patch region TM2, and route B cross the east of North Korea to the region TM 1.

## 4. Discussion

### 4.1. Suitable habitat

Water deer expansion into the transboundary region between the Russian Far East, northeast China, and North Korea is a recent event, and suitable habitat assessment provides insights into their expansion trends. Our study used current knowledge on this expansion combined with landscape factors to find suitable habitat in a larger region, including parts of the Korean Peninsula, northeast China, and the Russian Far East.

Our study predicted a suitable habitat in the east and west coasts of the Korean Peninsula, which was consistent with the information in the existing records. We analyzed the influence of landscape factors to find suitable habitats, which may reflect the current expansion in the range of water deer. Based on our results, beyond the current occurrence area, suitable habitat extends to the north in the Russian Far East, where there is lower human pressure as well as good water resources and vegetation. Water deer have high fecundity compared to other deer species, and each year female deer can give birth with three to four calves on average, with as maximum record of seven calves at once, and calves reach sexual maturity after one year [40]. With such a strong reproductive capacity, population numbers in the areas of expansion are likely to increase rapidly, and there is a high possibility that they will continue to expand their territory toward the north.

Low elevation, proximity to cropland and water, and presence of wetlands were important environmental factors according to our model. We believe that this may be related to water deer physiology and diet as well as competition with other deer species such as roe deer, but this assumption needs further research. Our findings in term of land use variable result from habitat model match with the knowledge of the species as GPS tracking data from water deer research in South Korea showed that forest, wetland, agriculture, and water areas were the most frequently used land covers [41]. Further, the preference for 20–25° slopes and broadleaved forests was significance related with water deer density [42]. Wetlands, water, and agricultural areas are landscape features of similar importance both in South Korea and the newly occupied region, although differences may arise due to varying geographical conditions in different areas. Maxent is suggested to be a highly accurate machine learning method to model species distribution when used for water deer habitat prediction in South Korea [25]. In our research, it also performed well for predicting habitat and influencing factors.

### 4.2. Dispersal routes

Based on the landscape connectivity analysis, two possible routes were estimated as having a high probability of water deer dispersal from the North Korean populations to the newly occupied region, and at the same time, no barriers could be observed between the two suitable areas in North Korea. The newly colonized region in the lower Tumen area was next to important protected areas, namely the Northeast China Tiger and Leopard National Park, the Khasansky provincial nature park and the Land of the Leopard National Park in Russia. In North Korea, protected areas may also play an important role in the dispersal of wildlife, such as the Suryong Mountain Animal Reserve (수룡산동물보후구) located in the Tosan county of North Hwanghae Province, which can connect the two suitable habitat patches region NK2 and region NK3. Along the predicted route B, protected areas such as Daeheung Animal Reserve (대흥동물보호구) and Donggye animal reserve (동계동물보호구) are present in Ryanggang Province, and Gwanmobong nature reserve (관모봉자연보호구) forest area in North Hamyeong province. Some protected areas for birds or fish may also contribute to the provision of suitable water resources and wetland habitat for water deer dispersal, such as Kuumya Migratory Bird Reserve (금야철새보호구) in South Hamgyong Province, and Rason Migratory Bird Reserve (라선철새보호구) in North Hamgeong Province, which connects the wetlands in the lower Tumen river with the wetlands in China and Russia. Animals’ dispersion through the route A may cross central northern North Korea into China and reach the newly occupied area by moving along the border. Within North Korea, Geumseok animal reserve (금석동물보호구) and Ogasan nature reserve (오가산자연보호구) may provide habitat for water deer dispersing northward across the Yalu river marking the China-North Korean border. From there, dispersing water deer can reach the forests in China in the Jilin Baishan Musk Deer National Nature Reserve (吉林白山原麝国家级自然保护区), or continue through the northern area of Changbai mountain to reach the Tiger and Leopard National Park, and subsequently reach the newly occupied region. The predicted route estimate is limited by the absence of data on animal movement, which may provide a better understanding of dispersal routes and habitat connectivity.

The water deer dispersal route may also provide insights for the conservation of big cats. The Changbai mountain region was predicted to be an important potential habitat for the recovery of an isolated Amur tiger population [43], and the connectivity between habitat patches is crucial for the success of their conservation. There is already evidence that amur tigers and leopards have started hunting this new prey species in the newly colonized region: unpublished camera trap photo of a leopard hunting water deer in China, and tiger scat with water deer teeth in Russia. Ecological research on tigers suggests the most important requirements for wild tiger survival are prey, water, and sheltered areas for hunting [44]. Prey availability is the most basic and important element for predators; thus, new prey habitats may contribute to the recovery of large cats in the landscape.

There are many barriers due to national security issues between countries in Northeast Asia, but the water deer expansion and our habitat connection analyses show that there is connectivity between sites for wildlife habitat in the region. Future surveys may have important implications for the conservation of the connected landscape.

### 4.3. Implications for conservation

Species based conservation in a single area is insufficient. Climate change and human activities drive species dispersal from one site to another, and a connected landscape for wildlife movement and survival reduces the risk for species facing threats in a single region, as well as mitigates pressure from environmental influence. Conservation planning needs to move from a single-site approach to one that can incorporate processes at landscape scales. In the case of water deer dispersal, a transboundary biodiversity conservation plan needs to be considered, as landscape connectivity in the whole region will contribute to the health of forest and wetland ecosystems.

Numerous issues can affect the success of landscape conservation, such as the lack of surveys, illegal hunting, unclear reserve boundaries, lack of awareness from local people, and insufficient collaboration between stakeholders. Further, there are many cases of water deer mortalities due to roadkill [45]; therefore, measures to prevent roadkill are also necessary. Based on the available information regarding water deer in South Korea, they consume crops, rice, and other agricultural products, and in areas with high quality habitat, different measures are needed to prevent conflicts between water deer and the local community [40]. Other direct threats, such as illegal hunting, need to be addressed by raising local community awareness and through projects mitigating conflicts between wildlife and humans. Knowledge exchange between different stakeholders within and between countries will not only contribute to the better management of one species but will also benefit the whole landscape and the ecosystem.

## 5. Conclusion

In our research, we analyzed water deer data from newly occupied areas in the transboundary region between China, Russia, and North Korea, and predicted suitable habitats and two possible dispersal routes connecting them. With this information, we provide a general understanding of the habitat use of this species, and demonstrate the existence of connectivity for wildlife in the larger landscape, which can contribute to the conservation of the whole landscape, although it will need more effort to survey and protect.

## Acknowledgements

The authors would like to thank Felipe Perez for their assistance with the English language review.

## Author Contributions

### Conceptualization

Ying LI, Hang LEE, Yongwon MO

### Field data collection

Ying LI, Hailong LI, Yury DARMAN, Gleb SEDASH

### Data preparation

Ying LI, Yuxi PENG, Hailong LI

### Data analysis and method

Ying LI, Yongwon MO, Dong Kun LEE, Tianming WANG

### Mapping

Ying LI, Yuxi PENG

### Funding acquisition

Ying LI,Weihong ZHU, Hang LEE

### Writing-original draft

Ying LI

### Writing-review and editing

Ying LI, Puneet PANDEY, Amaël BORZÉE

## Reference

1. Legagneux P, Gauthier G, Lecomte N, Schmidt NM, Reid D, Cadieux MC, et al. Arctic ecosystem structure and functioning shaped by climate and herbivore body size. Nature Climate Change. 2014;4(5):379–83. doi: 10.1038/nclimate2168.

2. Risch AC, Ochoa-Hueso R, van der Putten WH, Bump JK, Busse MD, Frey B, et al. Size-dependent loss of aboveground animals differentially affects grassland ecosystem coupling and functions. Nat Commun. 2018;9(1):3684. Epub 2018/09/13. doi: 10.1038/s41467-018-06105-4. PubMed PMID: 30206214; PubMed Central PMCID: PMCPMC6133970.

3. Hydropotes inermis. The IUCN Red list of Threatened Species 2015 [Internet]. 2015.

4. Jo Y, Baccus JT, Koprowski JL. Mammals of Korea: National Institute of Biological Resources; 2018.

5. Won Hg. Wildlife list in North Korea: Pyongyang book press; 1968.

6. MNA MNA. Protected Areas In Our Country Peony printing technician (모란인쇄기술사) 2005.

7. Darman YA, Storozhuk VB, Sedash GA. Hydropotes inermis (Cervidae), a new species for the Russian fauna registered in the Land of Leopard National Park (Russia). Nature Conservation Research. 2019;4(3):127–9. doi: 10.24189/ncr.2019.057.

8. Sheng H, Ohtaishi N, Houji L. The Mammalian of China: China Forestry Publishing House; 1999.

9. Jung J, Shimizu Y, Omasa K, Kim S, Lee S. Developing and testing a habitat suitability index model for Korean water deer (Hydropotes inermis argyropus) and its potential for landscape management decisions in Korea. Animal Cells and Systems. 2016;20(4):218–27. doi: 10.1080/19768354.2016.1210228.

10. Franklin J. Mapping species distributions: Cambridge University Press; 2009.

11. Hilts DJ, Belitz MW, Gehring TM, Pangle KL, Uzarski DG. Climate change and nutria range expansion in the Eastern United States. The Journal of Wildlife Management. 2019;83(3):591–8. doi: 10.1002/jwmg.21629.

12. Manish K, Pandit MK. Identifying conservation priorities for plant species in the Himalaya in current and future climates: A case study from Sikkim Himalaya, India. Biological Conservation. 2019;233:176–84. doi: 10.1016/j.biocon.2019.02.036.

13. Pearman-Gillman SB, Duveneck MJ, Murdoch JD, Donovan TM. Drivers and Consequences of Alternative Landscape Futures on Wildlife Distributions in New England, United States. Frontiers in Ecology and Evolution. 2020;8. doi: 10.3389/fevo.2020.00164.

14. Saito MU, Momose H, Inoue S, Kurashima O, Matsuda H. Range-expanding wildlife: modelling the distribution of large mammals in Japan, with management implications. International Journal of Geographical Information Science. 2014;30(1):20–35. doi: 10.1080/13658816.2014.952301.

15. Elith J, Graham CH, Anderson RP, Dudík M, Ferrier S, Guisan A, et al. Novel methods improve prediction of species’ distributions from occurrence data. ECOGRAPHY. 2006;29:129–51.

16. Xing D, Hao Z. The principle of maximum entropy and its applications in ecology. Biodiversity Science. 2011;19(3):295–302.

17. Evcin O, Kucuk O, Akturk E. Habitat suitability model with maximum entropy approach for European roe deer (Capreolus capreolus) in the Black Sea Region. Environ Monit Assess. 2019;191(11):669. Epub 2019/10/28. doi: 10.1007/s10661-019-7853-x. PubMed PMID: 31650357.

18. Dickson BG, Albano CM, Anantharaman R, Beier P, Fargione J, Graves TA, et al. Circuit-theory applications to connectivity science and conservation. Conserv Biol. 2019;33(2):239–49. Epub 2018/10/13. doi: 10.1111/cobi.13230. PubMed PMID: 30311266; PubMed Central PMCID: PMCPMC6727660.

19. Shah VB, McRae B. Circuitscape: A Tool for Landscape Ecology. Proceedings of the 7^th^Python in Science Conference 2008. p. 62–6.

20. Belyaev DA, Jo Y-S. Northernmost finding and further information on water deer Hydropotes inermis in Primorskiy Krai, Russia. Mammalia. 2020;85(1):71–3. doi: 10.1515/mammalia-2020-0008.

21. Li Z, Zhang Z, Mi S, Wu J, Xu T, Liu Z, et al. The complete mitochondrial genome of water deer in Liaoning, China. Mitochondrial DNA B Resour. 2020;5(1):922–3. Epub 2020/12/29. doi: 10.1080/23802359.2020.1719936. PubMed PMID: 33366811; PubMed Central PMCID: PMCPMC7748434.

22. Li Z-Z, Wu J-P, Teng L, Liu Z-S, Wang B-K, Liu Y-h, et al. The Rediscovery of Water Deer (Hydropotes Inermis) in Jilin Province. Chinese Journal of Zoology. 2019;54(1):108–12. doi: 10.13859/j.cjz.201901013.

23. GBIF Backbone Taxonomy [Internet]. 2021.

24. Aiello-Lammens ME, Boria RA, Radosavljevic A, Vilela B, Anderson RP. spThin: an R package for spatial thinning of species occurrence records for use in ecological niche models. Ecography. 2015;38(5):541–5. doi: 10.1111/ecog.01132.

25. Song W, Kim E. A Comparison of Machine Learning Species Distribution Methods for Habitat Analysis of the Korea Water Deer(Hydropotes inermis argyropus). Korean Journal of Remote Sensing. 2012;28(1):171–80.

26. Zhang E, Liwei T, Yongbei W. Habitat selection of the Chines water deer (Hydropotes inermis) in Yancheng Reserve, Jiangsu Province. Acta Theriologica Sinica. 2006;26(1):46–63.

27. Jarvis A, Reuter HI, Nelson A, Guevara E. Hole-filled seamless SRTM data V4, International Centre for Tropical Agriculture (CIAT) 2008. Available from: https://srtm.csi.cgiar.org.

28. Global land cover and land use 2019, v1.0 [Internet]. 2021. Available from: https://glad.umd.edu/dataset/global-land-cover-land-use-v1.

29. Openstreetmap [Internet]. 2015. Available from: https://planet.openstreetmap.org.

30. George D, Mallery P. IBM SPSS Statistics 26 Step by Step: A Simple Guide and Reference (6th ed.) 2019.

31. Kim B-J, Lee S-D. Home range study of the Korean water deer (Hydropotes inermis agyropus) using radio and GPS tracking in South Korea: comparison of daily and seasonal habitat use pattern. Journal of Ecology and Environment. 2011;34(4):365–70. doi: 10.5141/jefb.2011.038.

32. Elith J, Phillips SJ, Hastie T, Dudík M, Chee YE, Yates CJ. A statistical explanation of MaxEnt for ecologists. Diversity and Distributions. 2011;17(1):43–57. doi: 10.1111/j.1472-4642.2010.00725.x.

33. Phillips SJ, Anderson RP, Dudík M, Schapire RE, Blair ME. Opening the black box: an open-source release of Maxent. Ecography. 2017;40(7):887–93. doi: 10.1111/ecog.03049.

34. Zhu G, Qiao H. Effect of the Maxent model’s complexity on the prediction of species potential distributions. Biodiversity Science. 2016;24(10):8. doi: 10.17520/biods.2016265Biodiversity.

35. Cobos ME, Peterson AT, Barve N, Osorio-Olvera L. kuenm: an R package for detailed development of ecological niche models using Maxent. PeerJ. 2019;7:e6281. Epub 2019/02/14. doi: 10.7717/peerj.6281. PubMed PMID: 30755826; PubMed Central PMCID: PMCPMC6368831.

36. Mcrae BH, Dickson BG, Keitt TH, Shah VB. Using circuit theory to model connectivity in ecology, evolution, and conservation. Ecology. 2008;89(10):2712–24.

37. Feng H, Li Y, Li Y, Li N, Li Y, Hu Y, et al. Identifying and evaluating the ecological network of Siberian roe deer (Capreolus pygargus) in Tieli Forestry Bureau, northeast China. Global Ecology and Conservation. 2021;26. doi: 10.1016/j.gecco.2021.e01477.

38. Liu C, Newell G, White M, Bennett AF. Identifying wildlife corridors for the restoration of regional habitat connectivity: A multispecies approach and comparison of resistance surfaces. PLoS One. 2018;13(11):e0206071. Epub 2018/11/08. doi: 10.1371/journal.pone.0206071. PubMed PMID: 30403713; PubMed Central PMCID: PMCPMC6221308.

39. Brad MR, Viral S, Tanmay M. Circuitscape 4 User Guide: The Nature Conservancy; 2013. Available from: http://www.circuitscape.org.

40. Kim BJ. Korean Water Deer: National Institute of Ecology of South Korea; 2016.

41. Park H, Lee S. Habitat Use Pattern of Korean Waterdeer based on the Land Coverage Map. Journal of Wetlands Research. 2013;15(4):567–72.

42. Kim B-J, Oh D-H, Chun S-H, Lee S-D. Distribution, density, and habitat use of the Korean water deer (Hydropotes inermis argyropus) in Korea. Landscape and Ecological Engineering. 2010;7(2):291–7. doi: 10.1007/s11355-010-0127-y.

43. Hebblewhite M, Zimmermann F, Li Z, Miquelle DG, Zhang M, Sun H, et al. Is there a future for Amur tigers in a restored tiger conservation landscape in Northeast China? Animal Conservation. 2012;15(6):579–92. doi: 10.1111/j.1469-1795.2012.00552.x.

44. Schaller B. The Deer and the Tiger The university of Chicago Press, LTD; 1967.

45. Kim K, Yi Y, Woo D, Park T, Song E. Factors influencing Roadkill Hotspot in the Republic of Korea. PNIE. 2021;2(4):274–8. doi: 10.22920/PNIE.2021.2.4.274.

